# LFA-1 signals to promote actin polymerization and upstream migration in T cells

**DOI:** 10.1101/2020.04.29.069302

**Authors:** Nathan H Roy, Sarah Hyun Ji Kim, Alexander Buffone, Daniel Blumenthal, Bonnie Huang, Sangya Agarwal, Pamela L Schwartzberg, Daniel A Hammer, Janis K Burkhardt

**Affiliations:** Department of Pathology and Laboratory Medicine, Children’s Hospital of Philadelphia Research Institute, Philadelphia, PA, USA; Perelman School of Medicine, University of Pennsylvania, Philadelphia, PA, USA; Department of Bioengineering, University of Pennsylvania, Philadelphia, PA, USA; Department of Chemical and Biomolecular Engineering, University of Pennsylvania, Philadelphia, PA, USA; Laboratory of Immune System Biology, National Institute of Allergy and Infectious Diseases, Bethesda, Maryland.; National Human Genome Research Institute, National Institutes of Health, Bethesda, Maryland.

**Keywords:** T cell, integrins, migration, shear flow, signaling, actin

## Abstract

T cell entry into inflamed tissue requires firm adhesion, cell spreading, and migration along and through the endothelial wall. These events require the T cell integrins LFA-1 and VLA-4 and their endothelial ligands ICAM-1 and VCAM-1, respectively. T cells migrate against the direction of shear flow on ICAM-1 and with the direction of shear flow on VCAM-1, suggesting that these two ligands trigger distinct cellular responses. However, the contribution of specific signaling events downstream of LFA-1 and VLA-4 has not been explored. Using primary mouse T cells, we found that engagement of LFA-1, but not VLA-4, induces cell shape changes associated with rapid 2D migration. Moreover, LFA-1 ligation results in activation of the PI3K and ERK pathways, and phosphorylation of multiple kinases and adaptor proteins, while VLA-4 ligation triggers only a subset of these signaling events. Importantly, T cells lacking Crk adaptor proteins, key LFA-1 signaling intermediates, or the ubiquitin ligase cCbl, failed to migrate against the direction of shear flow on ICAM-1. These studies identify novel signaling differences downstream of LFA-1 and VLA-4 that drive T cell migratory behavior.

**Summary Statement:** Inflammatory responses require leukocyte migration along the vascular wall. We show that signaling from β2, but not β1, integrins induces cytoskeletal changes needed for upstream migration under shear flow.

## Introduction

Migration of leukocytes from the vasculature into peripheral tissue is central to their role in fighting pathogens, promoting tissue repair, and attacking solid tumors. This process, called transendothelial migration (TEM), is a key control point in the inflammatory response (1). TEM is a multi-step process that begins with selectin-dependent cell rolling on the vasculature, followed by chemokine-induced cell arrest. At this point, integrins expressed on the leukocyte interact with their endothelial ligands, resulting in shear resistant adhesion, cell spreading, and migration along the endothelium (1–3). Leukocyte migration along the vascular wall is a prerequisite for transmigration, and is thought to allow cells to search for ideal sites such as 3-way junctions to cross the endothelial layer (4–8). Intravital imaging of immune responses *in vivo* has revealed that leukocyte migration along the endothelial monolayer is not random and can be directed by shear flow forces (9, 10). Interestingly, leukocytes have been observed to preferentially migrate against the direction of shear flow (10), an unexpected result given the extra energy expenditure needed to oppose head-on shear forces. Although it is not clear why leukocytes display this phenotype, *in vitro* studies have shown that T cells crawling against the direction of shear on inflamed endothelia are more likely to undergo transmigration (11, 12), suggesting a link between these two mechanically demanding processes.

T cell adhesion and migration on the vascular wall are dependent on integrin interactions with their endothelial ligands. The two major integrin ligands expressed on inflamed endothelia are ICAM-1 and VCAM-1, which serve as binding partners for the T cell integrins LFA-1 (α_L_β_2_) and VLA-4 (α_4_β_1_), respectively. Although both of these integrin-ligand pairs can support adhesion, they seem to promote distinct migratory behaviors. The most striking example of this is observed under shear flow, in which T cells migrating on ICAM-1 coated surfaces preferentially migrate against the direction of shear flow (upstream migration), while T cells migrating on VCAM-1 coated surfaces migrate with the direction of shear flow (downstream migration) (13–17). This integrin-dependent phenomenon has also been documented in B cells (18), hematopoietic precursors (19), and neutrophils (under some conditions) (20). These data point towards fundamentally different roles for ICAM-1 and VCAM-1 in coordinating leukocyte migration.

A biophysical model has been proposed to explain the ability of T cells to migrate upstream on ICAM-1. The model is based on the finding that LFA-1/ICAM-1 interactions occur toward the front of the cell, and induce the formation of a broadly spread leading edge and raised, trailing uropod, whereas VLA-4/VCAM-1 interactions take place toward the back of the cell and fail to generate these shape changes (15). The shape changes induced by LFA-1 engagement allow the uropod to act as a “wind vane” that passively steers the T cell upstream (14). Although this model relies on the idea that localized integrin adhesion results in the relevant cell shape changes, T cell spreading and migration have also been shown to involve integrin signaling. This is best documented for LFA-1; T cells that come into contact with surface-presented ICAM-1 immediately polarize and begin to migrate (21–23).

Recently, we characterized a signaling pathway downstream of LFA-1 that leads to cell spreading, actin polymerization, and migration (24). Central players in this pathway are the Crk adaptor proteins, which coordinate Src-dependent phosphorylation of the scaffolding ubiquitin ligase c-Cbl, ultimately leading to PI3K activity and actin responses. Importantly, disruption of this pathway by deleting Crk proteins perturbs LFA-1 dependent migration, showing that LFA-1 signaling is an important factor driving T cell migration on ICAM-1. Because the LFA-1/ICAM-1 interaction triggers strong migratory responses in T cells, we hypothesized that differential integrin signaling events account for the differences in behavior of T cells migrating on ICAM-1 versus VCAM-1. Therefore, we analyzed T cell migration and signaling in response to binding ICAM-1 or VCAM-1, with the goal of determining how these different integrin ligands control T cell migration.

## Results

We and others have previously shown that T cells preferentially migrate against the direction of shear on ICAM-1 and with the direction of shear on VCAM-1 (11–17). To verify that our primary mouse T cells display this behavior, we activated T cells for two days and rested them for an additional three days in the presence of IL-2, a procedure that generates highly motile T cell blasts expressing both LFA-1 and VLA-4. These cells were allowed to adhere to surfaces coated with ICAM-1, VCAM-1, or both ligands mixed at a 1:1 ratio, and then subjected to shear flow of defined rates. Video microscopy was performed and the migrating cells were tracked and analyzed as described in the Materials and Methods. **Figure 1A** displays representative scattergrams of T cell migration tracks, with red tracks indicating upstream migration and blue tracks indicating downstream migration. These data show a clear tendency of T cells to migrate upstream on ICAM-1 and mixed surfaces, while migration on VCAM-1 was almost completely downstream. To quantify upstream versus downstream migration, we calculated the migration index (MI) for each cell. MI is determined by dividing the displacement in the X-direction (the axis parallel to the shear flow) by the total track length. An MI value of −1 indicates that cells migrate in a straight trajectory against the direction of flow, an MI value of +1 indicates migration in a straight trajectory with the direction of flow. Quantification from at least three independent experiments confirms that T cells tend to migrate against the direction of shear on ICAM-1 **(Fig 1B)**. T cells on mixed surfaces displayed a phenotype similar to those on ICAM-1 alone, showing that ICAM-1 ligation induces the dominant phenotype.

**Figure 1:**
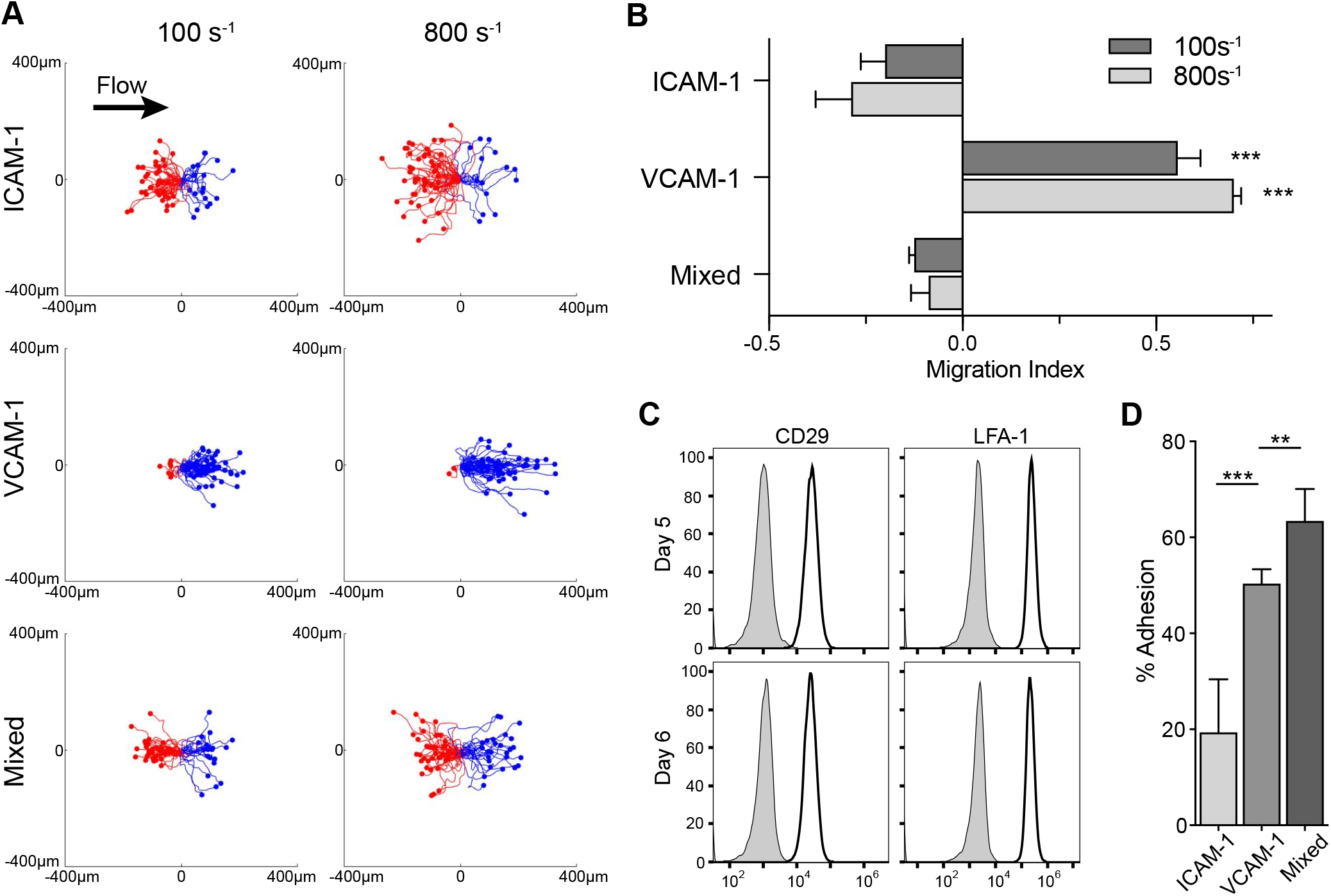
Activated primary mouse T cells migrate against the direction of shear on ICAM-1. **A)** Representative scattergrams of activated primary mouse CD4^+^ T cells migrating on ICAM-1, VCAM-1, or mixed surfaces under shear flow conditions (100s^−1^ or 800s^−1^). Red lines, cell tracks with a net migration against the direction of shear (upstream). Blue lines, cell tracks with a net migration in the direction of shear (downstream). **B)** Average migration index (MI) pooled from three independent experiments. MI for each cell equals the ratio of the displacement left or right by the total distance traveled. Migration with no directional preference is a value of 0. Negative values indicate net migration upstream, while positive values indicate downstream migration. **C)** Surface levels of CD29 (the β1 chain of VLA-4) and LFA-1 (αL/β2) on CD4^+^ T cells at day 5 and 6 post activation (the days used for all experiments). Gray shaded lines represent isotype controls. **D)** CD4^+^ T cell adhesion to the indicated ligands was measured using a plate based adhesion assay. Panels B and D show means +/− StDev for three independent experiments. Statistics were calculated using a one-way ANOVA, **p<0.01; ***p<0.001.

We reasoned that this upstream/downstream phenotype could result from large differences in LFA-1 or VLA-4 expression levels, or the ability of the different integrins to promote firm adhesion. To test this, we surface labeled primary T cells for LFA-1 and CD29 (the β1 subunit of VLA-4). T cells expressed appreciable surface levels of both integrins on days 5 and 6 after activation, the days that cells were used for experiments throughout this study **(Fig 1C)**. To compare the ability of the two integrins to support adhesion, we utilized a standard plate-based adhesion assay to measure binding to surfaces coated with ICAM-1, VCAM-1, or both ligands in combination. Interestingly, VCAM-1 supported more robust adhesion than ICAM-1, and the mixed surfaces promoted adhesion that was slightly higher than VCAM-1 alone **(Fig 1D)**. Thus, both integrins act as functional adhesion molecules, and there is no correlation between the efficiency of adhesion and upstream/downstream migratory behavior. Taken together, these data show that primary mouse T cells migrate upstream on ICAM-1 and downstream on VCAM-1, and that this phenotype cannot be explained simply by differences in integrin expression or adhesive functions.

To better understand the differential nature of migration on ICAM-1 and VCAM-1, we analyzed the shape and behavior of migrating T cells. This was done in the absence of shear to eliminate confounding effects induced by the shear itself. We showed previously that engagement of LFA-1 on primary mouse T cells by surface-bound ICAM-1 leads to cell spreading, cytoskeletal remodeling, and persistent directional migration (24). To ask if the VLA-4/VCAM-1 interaction induces similar responses, we conducted side-by-side analysis. Primary mouse T cells were allowed to interact with surfaces coated with ICAM-1, VCAM-1, or both ligands mixed at equal ratios. Poly-L-lysine was included as a non-specific control. Cells were then fixed, stained for F-actin and imaged using confocal microscopy. As shown in **Figure 2A**, T cells responded to ICAM-1 coated surfaces by spreading and forming a broad, actin-rich leading edge. In contrast, T cells responding to VCAM-1 displayed a more elongated shape. Although a leading edge could usually be identified, it was narrower and lacked robust F-actin accumulation. Quantification of F-actin levels confirmed that spreading on ICAM-1 induced an increase in actin polymer, whereas F-actin levels in cells responding to VCAM-1 remained unchanged relative to the poly-L-lysine control **(Fig 2B)**. T cells responding to mixed surfaces were indistinguishable from T cells responding to ICAM-1 alone **(Fig 2A-B)**, consistent with the finding that ICAM-1 induces the dominant phenotype. To evaluate how the different surfaces affect T cell migration, we tracked migrating cells with video microscopy. T cells migrating on ICAM-1 and mixed surfaces migrated significantly faster and more directionally than T cells on VCAM-1 **(Fig 2C-D)**. To more closely examine cell shape and actin dynamics during T cell migration, we utilized T cells expressing Lifeact-GFP. As observed in fixed cells labeled with phalloidin (**Fig 2A)**, T cells migrating on ICAM-1 or mixed surfaces adopted a spread morphology and displayed an actin rich leading edge, with a broad contact area with the substrate and a raised uropod **(Fig 2E-F)**. In contrast, T cells migrating on VCAM-1 showed less contact with the stimulatory surface and seem to adhere mostly through the center and uropod of the cell **(Fig 2E-F)**. This morphology is consistent with observations by others (14, 15). Analysis of the movies revealed that while T cells migrating on ICAM-1 coated surfaces moved smoothly, showing sustained actin polymerization at the leading edge, T cells on VCAM-1 had periodic bursts of movement concomitant with brief actin flares at the front of the cell **(Fig 2F-G**, see Supplemental Movies 1 and 2). This differential behavior can be readily observed using a kymographic representation **(Fig 2G)**. Altogether, these data show that engagement of ICAM-1, but not VCAM-1, results in T cell spreading, actin polymerization, and fast migration. Additionally, these data suggest that ICAM-1 triggers specific signaling events that lead to these responses.

**Figure 2:**
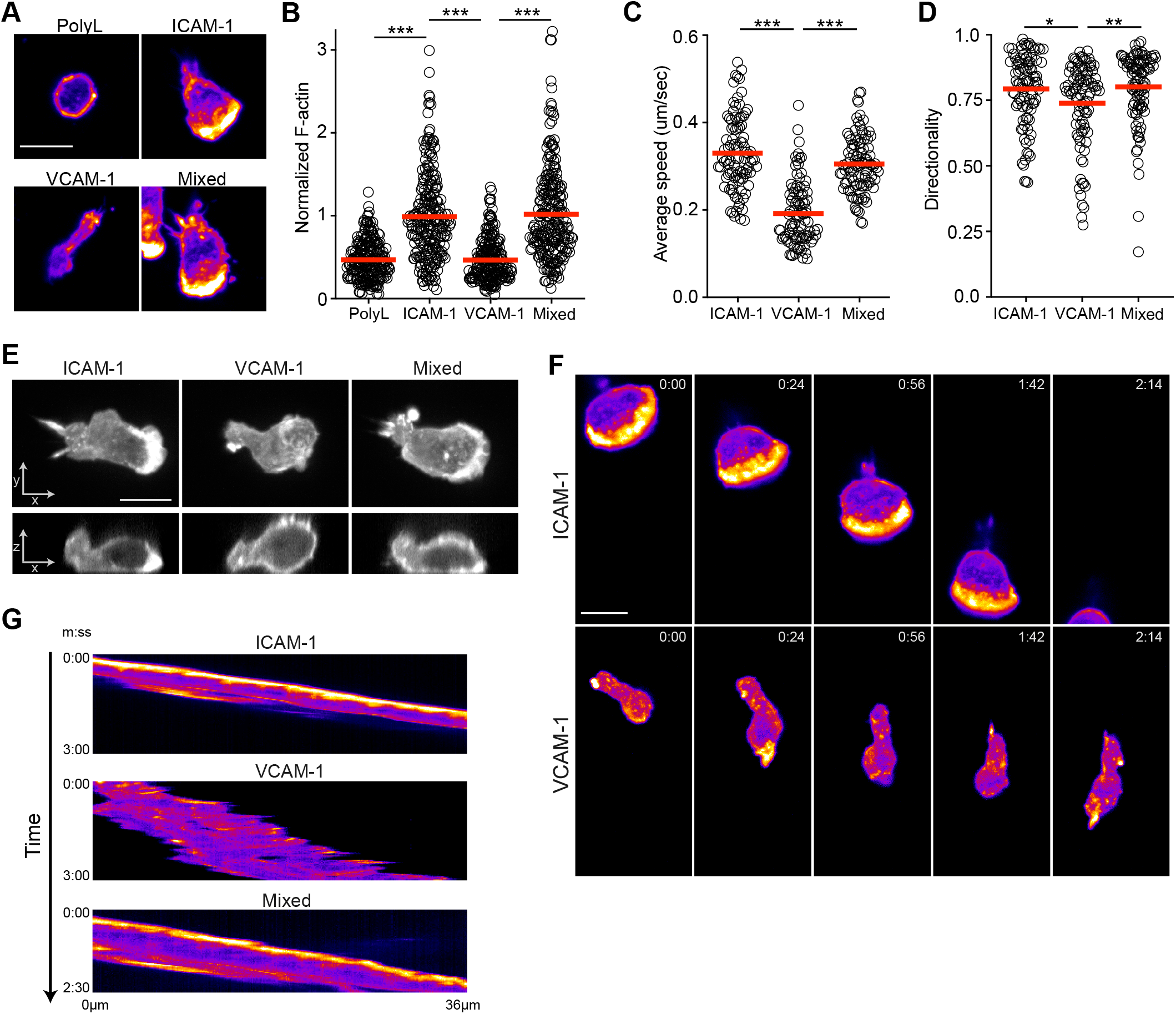
Quantitative and qualitative differences in T cell migration on ICAM-1 and VCAM-1. **A)** CD4^+^ T cells were allowed to migrate on the indicated ligands under static conditions, fixed, and stained with phalloidin to visualize F-actin. Images are a projection of four z-slices totaling 0.75μm, displayed as heat maps based on pixel intensity. Scale bar = 10μm. **B)** Quantification of total F-actin intensity per cell (each circle represents one cell). **C-D)** T cells were tracked while migrating on the indicated surfaces in the absence of shear, and the **(C)** average speed and **(D)** directionality were calculated. **E)** Orthogonal views of T cells expressing Lifeact-GFP, migrating on surfaces coated with the indicated ligands. Scale bar = 10μm. **F)** Migrating T cells expressing Lifeact-GFP were imaged using time-lapse confocal microscopy. Images are a projection of a 2μm total stack starting at the cell-coverslip interface. Scale bar = 10μm. See Supplemental Movies 1 and 2. **G)** Kymographs of T cells migrating as in **F**. Data in B-D are pooled from three independent experiments. Statistics were calculated using a one-way ANOVA, *p<0.05; **p<0.01; ***p<0.001.

Recently, we identified the Crk family adaptor proteins as key signaling intermediates that promote actin polymerization and migration in T cells downstream of LFA-1 (24). The Crk family consists of three proteins, Crk I, Crk II, and CrkL, which are transcribed from two loci (Crk I and Crk II are splice variants). In the absence of shear flow, T cells lacking all three Crk proteins (DKO T cells) migrate significantly slower than WT T cells on ICAM-1, and fail to spread and polymerize actin at the leading edge ((24) and **Fig 3A-B**). This phenotype is strikingly similar to that of WT T cells migrating on VCAM-1, suggesting that Crk protein signaling may be a determining factor in the cellular response to ICAM-1. To ask if Crk signaling is also important for upstream migration, we imaged WT and DKO T cells migrating in the presence of shear flow as in Figure 1. We found that DKO T cells were unable to migrate upstream on ICAM-1, with no apparent defect in downstream migration on VCAM-1 (**Fig 3C-D**). These data support a model in which Crk dependent LFA-1 signaling drives T cell spreading, actin responses, and upstream migration.

**Figure 3:**
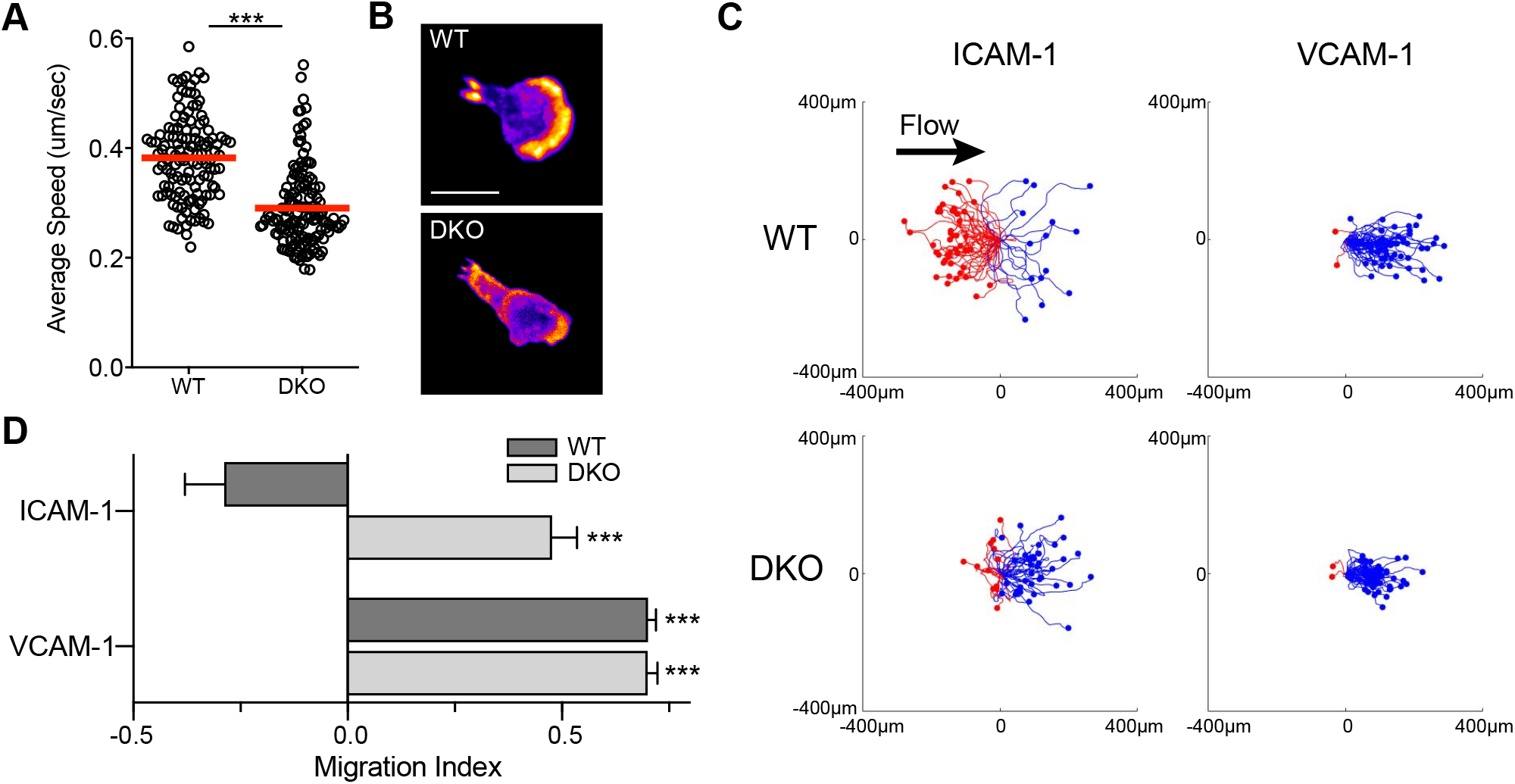
Crk deficient T cells fail to migrate upstream on ICAM-1. **A-B)** WT and DKO T cells were imaged while migrating on ICAM-1 in the absence of shear. **A)** Average speed pooled from three independent experiments. **B)** Representative images of migrating T cells expressing Lifeact-GFP, displayed as in Fig. 2. Scale bar = 10μm. **C)** Representative scattergrams of WT and DKO T cells imaged while migrating on ICAM-1 or VCAM-1 under shear flow (shear rate 800s^−1^). Red lines, cell tracks with a net migration against the direction of shear (upstream). Blue lines, cell tracks with a net migration in the direction of shear (downstream). **D)** Average migration index calculated from three independent experiments. Displayed as means +/− StDev. Statistics were calculated using a one-way ANOVA, *p<0.05; **p<0.01; ***p<0.001.

We next sought to identify signaling differences downstream of LFA-1 and VLA-4 that could account for the differential cell behavior that we observe. In addition to activating canonical signaling pathways, e.g. PI3K/AKT and ERK, engagement of LFA-1 and VLA-4 induces the phosphorylation and activation of several signaling and scaffold proteins, including CasL, cCbl, FAK, and Pyk2 (25–32). Crk proteins are crucial for multiple signaling events downstream of LFA-1, including phosphorylation of CasL and cCbl, as well as activation of the PI3K pathway (24). Our observation that Crk-deficient T cells do not migrate upstream on ICAM-1 suggests that some of these signaling events may be necessary for upstream migration. If so, we reasoned that these events might occur in cells responding to ICAM-1, but not to VCAM-1. To our knowledge, a direct comparison of signaling events initiated by surface-presented ICAM-1 versus VCAM-1 has never been reported. To determine if ICAM-1 and VCAM-1 differentially activate any of these downstream signaling pathways, we allowed T cells to interact with surfaces coated with either ICAM-1 or VCAM-1. The T cells were then lysed, and the activation state of the PI3K and ERK pathways were probed using phospho-specific antibodies. In addition, lysates were immunoprecipitated with anti-phosphotyrosine, and probed with antibodies to CasL, cCbl, and Pyk2. ICAM-1 stimulated robust phosphorylation of AKT and ERK, while stimulation with VCAM-1 induced only modest activation of these pathways **(Fig 4A-B)**. Similarly, robust phosphorylation of cCbl was only induced by ICAM-1 **(Fig 4A-B)**. Importantly, phosphorylation of CasL and Pyk2 occurred with similar efficiency after engagement of ICAM-1 and VCAM-1. These data demonstrate that both LFA-1 and VLA-4 are capable of signaling, but the repertoire of downstream signaling events is distinct.

**Figure 4:**
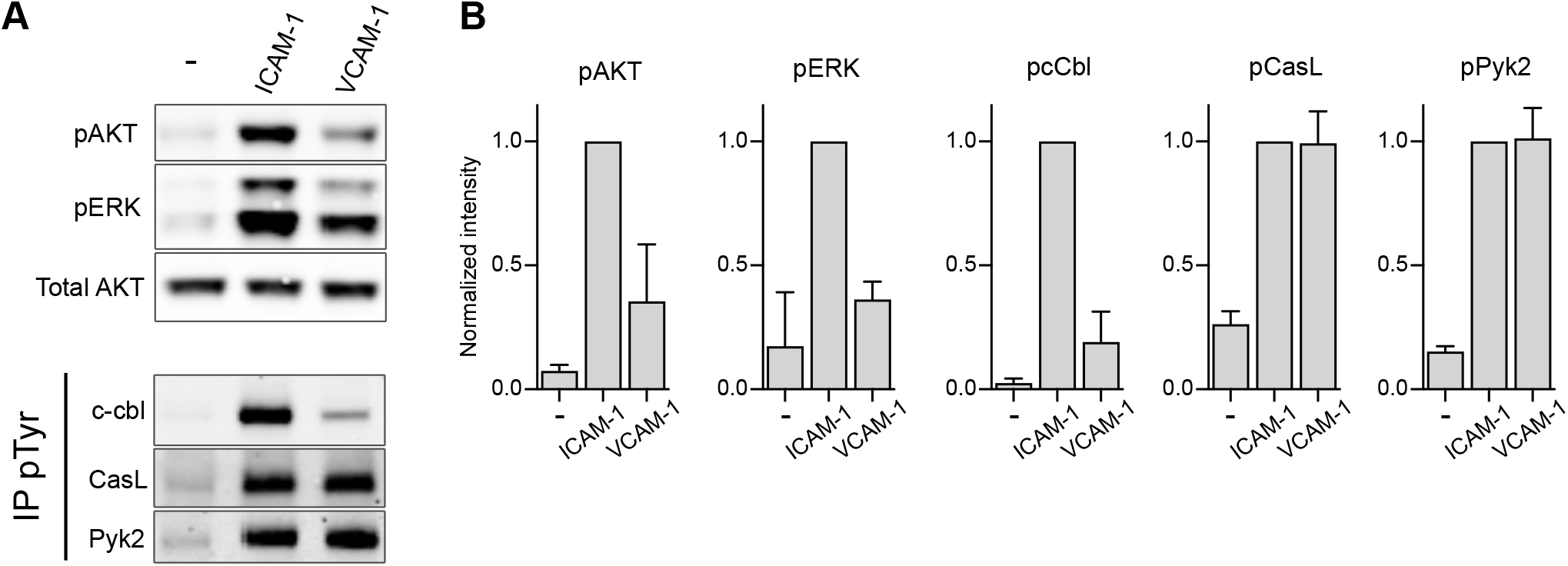
Differential signaling induced by ICAM-1 and VCAM-1. **A)** CD4^+^ T cells were allowed to settle on ICAM-1 or VCAM-1 coated plates for 20 mins and lysed. Lysates were immunoblotted with the indicated antibodies *(top panel)*, or immunoprecipitated with anti-pTyr, and immunoblotted with indicated antibodies *(bottom panel)*. **B)** Quantification of immunoblots performed as in A. Data represent means +/− StDev, normalized to the non-targeting control (NT), from three independent experiments.

Due to the clear differences in LFA-1 and VLA-4 signaling, we next wanted to directly test if any of the LFA-1 specific signaling events were important for upstream migration. In particular, we focused on cCbl and PI3K, both of which show defective activation in Crk-deficient T cells. To target these proteins, we implemented a CRISPR KO system in primary mouse T cells (33). T cell blasts cultured from Cas9-expressing mice were transduced with retroviral vectors containing non-targeting (NT), cCbl, or PI3Kδ gRNAs. T cells were then selected in puromycin for 3 days and used for experiments. Protein reduction was confirmed by immunoblotting **(Fig 5A-B)**. To assess the role of these proteins in LFA-1 mediated responses under static conditions, we allowed T cells to interact with ICAM-1 coated surfaces and stained for F-actin. T cells expressing NT gRNA showed the canonical migratory phenotype, with a broad actin-rich leading edge **(Fig 5C)**. Deletion of cCbl had no effect on this phenotype, while loss of PI3Kδ moderately reduced both cell spreading and actin polymerization **(Fig 5C-D)**. To test responses under shear flow conditions, we tracked cells migrating on ICAM-1 coated surfaces with a shear rate of 800s^−1^ and calculated MI. Interestingly, T cells expressing either NT or the PI3Kδ gRNA remained capable of migrating upstream on ICAM-1, while T cells expressing the cCbl gRNA failed to migrate upstream **(Fig 5E)**. This pattern also held true when analyzing the percent of cells migrating upstream **(Fig 5F)**. To best visualize this phenotype, we plotted the number of cells as a function of their MI for all the cells analyzed in **Figure 5F (Fig 5G)**. When compared to control NT T cells, a large proportion of T cells lacking cCbl had an MI between 0.1 and 0.4. This was also seen in representative scattergrams **(Supp Fig 1)**. The finding that T cells expressing the PI3Kδ gRNA showed normal upstream motility was rather surprising, especially since these cells did exhibit diminished spreading and polarization under static conditions. We reasoned that this might be because deletion of PI3Kδ was incomplete. Alternatively, other isoforms of PI3K could play a role. To address these possibilities, we repeated this analysis using T cells treated with the pan-PI3K inhibitor LY294002. As shown in **Supp Fig 2**, this inhibitor had no effect on upstream migration. This finding is consistent with previous work in human T cells (14, 17). Taken together, these studies identify cCbl, but not PI3K, as a necessary component of the LFA-1 signaling pathway that drives upstream migration.

**Figure 5:**
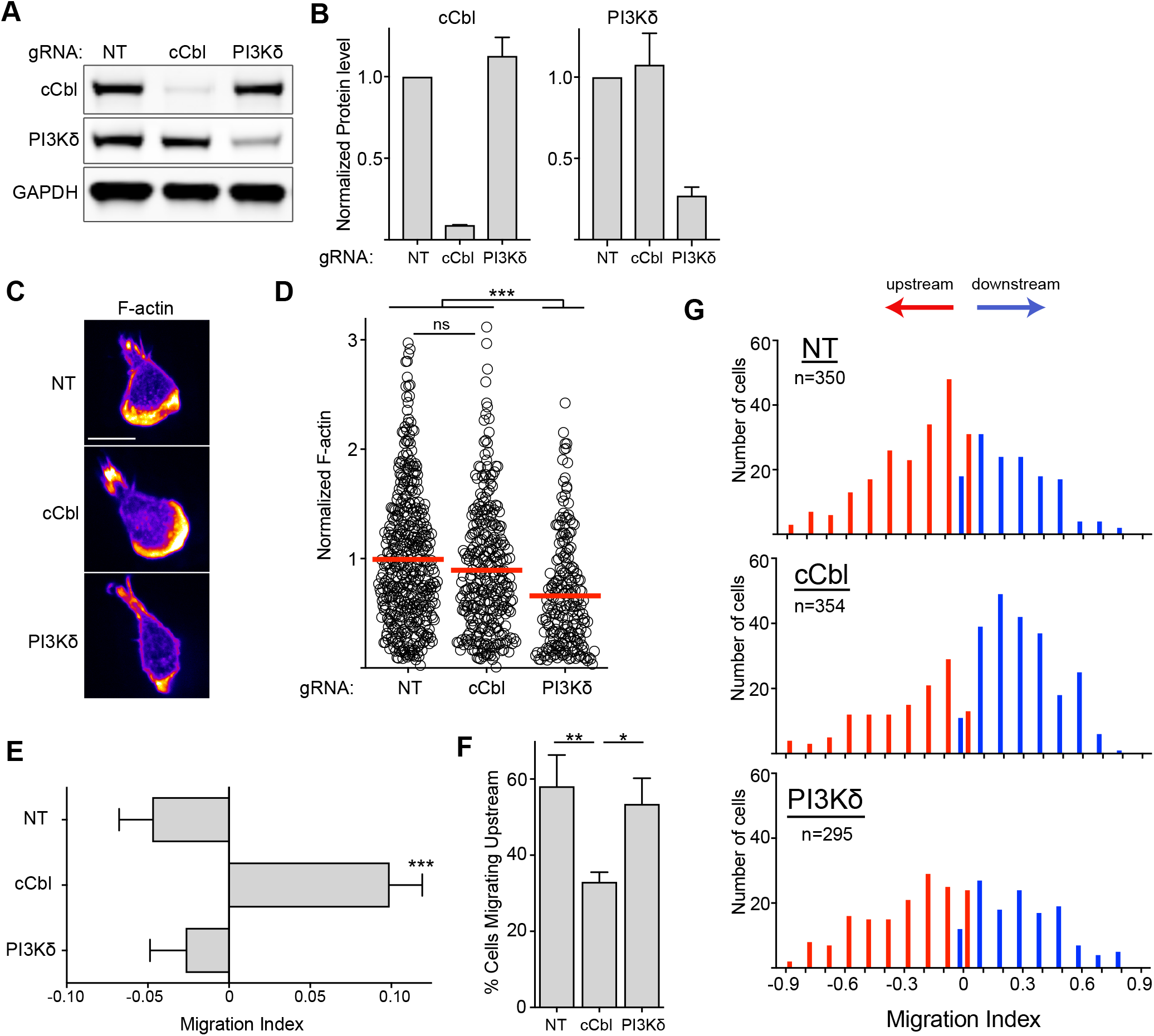
cCbl knockout reverses upstream migration on ICAM-1. **A)** CD4^+^ T cells from Cas9 expressing mice were transduced with the indicated gRNAs, selected for 3 days in puromycin, and then lysed and immunoblotted for the indicated proteins. **B)** Quantification of immunoblots performed as in A. Data represent means +/− StDev, normalized to the non-targeting control (NT), from three independent experiments. **C-D)** T cells expressing the indicated gRNAs were allowed to migrate on ICAM-1 coated surfaces, fixed, and stained with fluorescent phalloidin. **C)** Representative images, displayed as in Fig. 2. Scale bar = 10μm. **D)** Quantification of F-actin intensity per cell, pooled from three independent experiments. **E)** Migration index of T cells expressing the indicated gRNAs migrating on ICAM-1 under shear flow (shear rate 800s^−1^). **F)** Percent cells migrating upstream from experiments in panel E. **G)** Distribution of cells with a given MI pooled from 3 independent experiments. Panels E and F show means +/− SEM for three independent experiments. Statistics were calculated using a one-way ANOVA, *p<0.05; **p<0.01; ***p<0.001.

## Discussion

We and others have previously shown that T cells migrate upstream on ICAM-1 and downstream on VCAM-1 (11–17). In this study, we investigated the role of integrin signaling events in driving upstream migration. We found that engagement of LFA-1 by surface bound ICAM-1 triggered robust activation of many signaling events as well as cytoskeletal remodeling and cell shape changes, while engagement of VLA-4 by VCAM-1 failed to trigger some, but not all, of these events. Importantly, upstream migration on ICAM-1 could be reversed by deleting key proteins associated with LFA-1 signaling. Taken together, our data uncover fundamental signaling differences between LFA-1 and VLA-4 that directly affect T cell migration under shear flow.

Our first indication that LFA-1 signaling drives T cell migration came from our previous studies on the Crk adaptor proteins. We found that T cells lacking Crk proteins have defects in LFA-1 signaling cascades leading to activation of the PI3K pathway and phosphorylation of the ubiquitin ligase/scaffold protein cCbl. These defects are associated with impaired T cell spreading, actin polymerization, and migration in response to surface-bound ICAM-1 under static conditions (24). Here, we show that Crk deficient T cells fail to migrate against the direction of shear flow on ICAM-1, suggesting a role for signaling in upstream migration. Importantly, while WT T cells responding to ICAM-1 adopt the canonical, flattened morphology with a low-profile leading edge and a raised uropod (21–24, 31), DKO T cells fail to spread; they sit high on the coverslip even at the front. Thus, even when interacting with ICAM-1, DKO T cells look strikingly similar to WT T cells migrating on VCAM-1.

The phenotype of DKO T cells, which fail to flatten and migrate upstream, fits well with the passive steering mechanism proposed by the Valignat and co-workers, who argue that the direction of migration under shear is dictated by the shear forces acting on cells of different shape (14, 15). Central to this model is the observation that ICAM-1 induces a spread morphology, with the exception of the aforementioned uropod sticking up in the rear. Shear forces over the spread cell make the uropod act as a “wind vane”, aligning the cell to face against the direction of shear. On the other hand, cells migrating on VCAM-1 adhere mostly in the back, resulting in a raised front. In this instance, shear forces strike the front of the cell, pushing it downstream. In its simplest form, the passive steering mechanism does not explicitly require integrin signaling. Indeed, a signal-independent mechanism was proposed based on the lack of Ca^2+^ influx in cells migrating on either ICAM-1 or VCAM-1 (15). However, although it is clear that Ca^2+^ is important for neutrophil migration (34–36), a requirement for Ca^2+^ signaling in integrin-dependent T cell migration is not well established. Additionally, the lack of Ca2+ signaling does not rule out a role for other signaling events. Numerous kinases and adapter proteins are phosphorylated downstream of integrin engagement in T cells (37), and at least some of these events depend on Crk protein expression. Thus, we conclude that Crk proteins function as signaling intermediates in the pathway linking LFA-1 engagement and upstream motility. Importantly, signaling and passive steering are not mutually exclusive. Indeed, we believe that the two are inextricably linked. We postulate a two-step mechanism in which Crk-dependent signaling downstream of LFA-1 drives cell shape changes, thereby allowing the cell to undergo passive steering.

To further explore the role of signaling in T cell migration under shear flow, we compared signaling events downstream of LFA-1 versus VLA-4 engagement. We found that engagement of LFA-1 by surface bound ICAM-1 triggers robust signaling, including the activation of the PI3K and ERK pathways, as well as the phosphorylation and activation of scaffold and signaling proteins such as cCbl, CasL, and Pyk2. In contrast, engagement of VLA-4 by VCAM-1 does not trigger substantial activation of PI3K or ERK, nor phosphorylation of cCbl. We were particularly interested in the LFA-1 dependent activation of PI3K and cCbl, as these events are also blunted in Crk deficient T cells (24). Knockout of PI3Kδ or pharmacological inhibition of PI3K had no effect on the ability of T cell to migrate upstream, consistent with earlier studies (14, 17). However, knockout of cCbl did perturb upstream migration. Curiously, however, cCbl KO T cells looked morphologically indistinguishable from WT T cells on ICAM-1. This represents an interesting exception to the idea that signaling works by inducing cell shape changes that allow passive steering, and indicates that cCbl promotes upstream migration through a distinct mechanism.

How does cCbl control T cell migration under shear flow? We showed previously that cCbl interacts with the p85 subunit of PI3K after LFA-1 ligation (24), but this does not seem to contribute to upstream migration, as inhibition of PI3K activity had no effect. In addition to p85, cCbl is known to interact with dozens of different proteins in a cell-type dependent manner (38). Thus, one or more other binding partners could affect T cell migration. In addition to its scaffold function, cCbl functions as a RING finger ubiquitin ligase. Indeed, most of the work in T cells has focused on the ubiquitin ligase activity of cCbl, and it is generally accepted that cCbl acts as a negative regulator of T cell receptor signaling (39–44). The role of cCbl dependent ubiquitylation in T cell migration is largely unexplored. Since ubiquitylation can control protein degradation, trafficking, or functioning, it is attractive to speculate that cCbl promotes upstream migration under shear flow conditions by ubiquitylating proteins needed for T cell migration. Future work aimed at determining the molecular mechanism of cCbl function will provide valuable new insights into T cell migration.

Taken together, our findings demonstrate a clear role for integrin-dependent signaling during T cell migration, and reveal that LFA-1 and VLA-4 deliver distinct signals that direct different migratory behaviors under both static and shear-flow conditions. Since many vessels express multiple integrin ligands, and ligand levels change with tissue type and inflammatory status, it will be important going forward to understand how T cells integrate integrin signaling pathways. Indeed, integrin crosstalk has already been shown to play a role during upstream migration (17, 20). It will also be important to test how differential integrin signaling influences immune cell trafficking *in vivo*, especially during infiltration into sites of inflammation and solid tumors. A greater understanding of these events could guide the rational design of therapeutics to alter immune cell migration.

## Materials and Methods

### Antibodies and Reagents

Anti-CD3 clone 2C-11 and the anti-CD28 clone PV1 were obtained from BioXCell. Anti-pTyr clone PY-20 was from Upsate (Millipore). Anti-HEF1 (CasL) clone 2G9 and anti-Pyk2 clone YE353 were obtained from Abcam. Anti-pERK (9101), anti-pAKT (4060), and anti-c-Cbl (2747) were from Cell Signaling. Anti-AKT (559028) was from BD. Secondary antibodies conjugated to appropriate fluorophores and AlexaFluor-conjugated phalloidin were obtained from ThermoFisher Scientific. Recombinant mouse ICAM-1-Fc and VCAM-1-Fc were purchased from R&D Systems. Antibodies for flow cytometery were from BioLegend. These included rat anti-CD4 APC (clone RM4-5), rat anti-CD11a/CD18 (LFA-1) PE (clone H155-78), Armenian hamster anti-CD29 FITC (clone HMβ1-1), anti-Armenian hamster IgG (clone HTK888) FITC, and rat anti-Rat IgG1κ (clone RTK2071).

### Mice and T Cell Culture

All mice were housed in the Children’s Hospital of Philadelphia animal facility, according to guidelines put forth by the Institutional Animal Care and Use Committee. C57BL/6 mice, originally obtained from The Jackson Laboratory, were used as a source of WT T cells. Mice expressing Lifeact-GFP have been described previously (45). Additionally, mice lacking the Crk adaptor proteins in T cells (herein referred to as DKO) were generated by crossing CD4^+^ Cre mice with mice in which the two Crk loci have been floxed (Crk fl/fl:CrkL fl/fl mice) (46, 47). To generate DKO mice expressing Lifeact-GFP, Crk fl/fl:CrkL fl/fl mice were crossed with Lifeact-GFP mice, and the resulting Lifeact-GFP Crk fl/fl:CrkL fl/fl mice were crossed with CD4^+^ Cre mice. Mice expressing Cas9 (C57BL/6J-congenic H11^Cas9^, Jackson stock no. 028239) were used for CRISPR studies.

Primary mouse CD4^+^ T cells were purified from lymph nodes and spleens by negative selection. Briefly, after removing red blood cells by ACK lysis, cells were washed and incubated with anti-MHCII and anti-CD8 hybridoma supernatants (M5/114 and 2.43, respectively) for 20 min at 4°C. After washing, cells were mixed with anti-rat Ig magnetic beads (Qiagen BioMag), incubated for 15 min at 4°C, and subjected to three rounds of magnetic separation using a bench top magnet. The resulting CD4^+^ T cells were then immediately activated on 24-well plates coated with anti-CD3 and anti-CD28 (2C11 and PV1, 1 μg/ml each) at 1×10^6^ cells per well. Activation was done in T cell complete media, composed of DMEM (Gibco 11885-084) supplemented with 5% FBS, penicillin/streptomycin, non-essential amino acids, Glutamax, and 2μl 2-mercaptoethanol. Unless otherwise indicated, all tissue culture reagents were from Gibco. After 48h, cells were removed from activation and mixed at a 1:1 volume ratio with complete T cell media containing recombinant IL-2 (obtained through the NIH AIDS Reagent Program, Division of AIDS, NIAID, NIH from Dr. Maurice Gately, Hoffmann - La Roche Inc), to give a final IL-2 concentration of 20 units/mL. T cells were used at day 5-6 after activation.

### Flow Cytometry

Cells were first stained with Live/Dead aqua (ThermoFisher) in PBS following the manufacturer’s protocol. Staining was quenched using 1% bovine serum albumin solution. For staining, antibodies were diluted 1:100 in FACS buffer (PBS, 5% FBS, 0.02% NaN_3_, and 1 mM EDTA). Flow cytometry was performed using either a Cytoflex LX or CytoFlex S cytometer (Beckman Coulter) and data was analyzed using FlowJo software (FlowJo LLC). T cells were gated based on size, live cells, and expression of CD4^+^.

### Static Adhesion Assay

96 well plates (MaxiSorp, ThermoFisher) were coated with 2 μg/mL mouse ICAM-1, VCAM-1, or both ligands mixed (1 μg/mL each), in PBS overnight at 4°C. Plates were then washed 3x with PBS, blocked with 1% BSA in PBS for 1h at RT, and washed twice with PBS. Activated CD4^+^ T cells were used on day 5 after initial isolation. To prepare the cells, they were first labeled with Calcien AM (ThermoFisher) at a final concentration of 2.5 μM for 30 min at 37°C in serum-free DMEM. Cells were then washed and resuspended in T cell complete media, and incubated at 37°C for 30 min. Cells were washed and resuspended in 2.5% BSA in PBS (with Ca^2+^ and Mg^2+^) and 1×10^5^ cells were added to each well on ice in triplicate. After a 20 min incubation, the plate was read on a Bio-Tek Synergy HT fluorescence plate reader to obtain the baseline measurements representing “maximum adhesion” per well. The plate was then incubated at 37°C for 10 min, washed, and read again. The plate was washed and read a total of 2-4 times, until the signal from the unstimulated control was stable. To calculate the percent adhesion, the fluorescence per well after washes was divided by the initial “maximum adhesion” fluorescence reading per well. Background was subtracted using values obtained from empty wells.

### F-actin Quantification in migrating T cells

Lab-Tek 8 chamber slides (ThermoFisher) were coated overnight at 4°C with 2 μg/mL ICAM-1, VCAM-1, or both ligands mixed (1 μg/mL each). Activated CD4^+^ T cells were resuspended in Leibovitz’s L-15 media (Gibco) supplemented with 2 mg/mL glucose and incubated at 37°C for 20 min. T cells were then added to chamber slides for 20 min at 37°C followed by fixation in 3.7% paraformaldehyde in PBS. Cells were then blocked and permeabilized in PSG (PBS, 0.01% saponin, 0.05% fish skin gelatin) for 20 min, followed by 45 min with fluorescent phalloidin (Molecular Probes) in PSG. Cells were washed in PSG, mounted, and imaged using a 63x PlanApo 1.4 NA objective on an Axiovert 200M (Zeiss) with a spinning disk confocal system (Ultraview ERS6; PerkinElmer). Four z-planes spanning a total of 0.75 μm were collected at the cell-surface interface with an Orca Flash 4.0 camera (Hamamatsu). Image analysis of the rendered stacks was conducted using Velocity v6.3 software. Cells were identified using the “Find Objects” command, using a low threshold on the actin channel, and total phalloidin staining was quantified per cell based on integrated pixel intensity.

For the visualization of F-actin in live cells, Lifeact-GFP mice served as the source of CD4^+^ T cells. Activated CD4^+^ T cells were added to Lab-Tek 8 chamber slides for 20mins, gently washed in warm L-15 media, mounted, and imaged at 37°C using the spinning disk confocal system. For migration movies, 9 z-planes spanning a total of 2 μm were collected every 2 seconds. Projections of the full 2 μm were prepared in Fiji. For whole cell reconstruction, 49 z-planes spanning a total of 12 μm were captured, and rendered using the orthogonal viewing tool in Fiji.

### Migration Under Shear Flow

Surfaces were prepared by stamping 2 μg/ml of protein A/G (Biovision, San Francisco, CA) using PDMS stamps, onto UV ozone-treated, PDMS spin-coated glass slides. Surfaces were then blocked with 0.2% Pluronic F-127 (Sigma), washed, and subsequently incubated with 2 μg/ml of murine ICAM-1 Fc, VCAM-1 Fc, or a 1:1 mixture at 4°C overnight. Activated CD4^+^ T cells were resuspended in RPMI-1640 supplemented with 2 mg/ml D-glucose and 0.1% BSA and introduced into a flow chamber using a syringe. T cells were allowed to adhere for 30 min prior to the onset of shear (rate 100s^−1^ or 800s^−1^). Cells were then imaged every minute for 30 min using a Nikon TE300 with a custom built environmental chamber at 37°C and 5% CO_2_. The movies were exported to ImageJ and cells were tracked and analyzed using the manual tracking plugin in ImageJ and a custom MATLAB script. Only cells that stayed in the field-of-view the whole movie were included in the analysis. Migration Index (MI) describes the directionality of a cell by quantifying a ratio of a cell’s axial displacement to total distance it has traveled. With flow from left to right of field-of-view, negative MI represents cells traveling upstream, while positive MI denotes cells traveling downstream.

### Migration in Static Conditions

Lab-Tek 8 chamber slides (ThermoFisher) were coated with 2 μg/mL ICAM-1, VCAM-1, or both ligands mixed, overnight at 4°C. Activated CD4^+^ T cells were washed and resuspended in L-15 media containing 2 mg/mL D-glucose. T cells were then added to the chambers, incubated 20 min, gently washed to remove all unbound cells, and imaged using a 10x phase contrast objective at 37°C on a Zeiss Axiovert 200M microscope equipped with an automated X-Y stage and a Roper EMCCD camera. Time lapse images were collected every 30 sec for 10 min using SlideBook 6 software (Intelligent Imaging Innovations). Movies were exported into ImageJ, and cells were tracked using the manual tracking plugin to calculate speed. Directionality was calculated by take the ratio of displacement by track length.

### Retroviral Production

The recently described gRNA retroviral transfer vector MRIG (33) was modified to express the puromycin resistance gene in place of GFP. Specific gRNAs were cloned into this vector exactly as described in (33). The gRNA sequences used in this study were as follows, NT: 5’ GCGAGGTATTCGGCTCCGCG, cCbl: 5’ TGTCCCTTCTAGCCGCCCAG, and PI3Kδ: 5’ GGAGCGTGGGCGCATCACGG. 293T cells were transfected with MRIG-puro along with the retroviral packaging vector pCL-eco at a 4:3 ratio using the calcium phosphate method and allowed to incubate overnight. The next morning, the media was gently replaced, and cells were incubated for 24-30 hours. Viral-containing supernatants were harvested and centrifuged at 1000g for 10 min to remove cellular debris. Polybrene was added to clarified viral supernatants to a final concentration of 8 μg/mL, and supernatants were used immediately for T cell transductions.

### T cell transductions

CD4^+^ T cells harvested from Cas9 expressing mice were activated on 24-well plates coated with anti-CD3 and anti-CD28 (2C11 5 μg/ml and PV1 2 μg/ml) at 1×10^6^ cells per well in T cell complete media. After 24 hours, the conditioned media was removed and reserved for later use, and gently replaced with viral supernatants (with polybrene), and incubated at 37°C for 10 mins. The plate was then centrifuged at 1100g for 2h at 37°C. Directly following spinocculation, the plate was placed back in the incubator for 10 min. Viral supernatants were then removed and replaced with a mixture of fresh complete T cell media mixed with conditioned media at a 3:1 volume ratio, after which cells were cultured for an additional 24 hours. T cells were then removed from activation, and mixed at a 1:1 volume ratio with complete T cell media containing recombinant IL-2 to yield a final IL-2 concentration of 20 units/mL. After 4 hours, puromycin was added to a final concentration of 4 μg/mL. Cells were used after 3 days of selection (day 5 after isolation).

### Biochemical analysis of cell signaling in response to surface-bound ICAM-1 or VCAM-1

60mm tissue culture dishes (Corning, 430166) were coated with 2 μg/mL ICAM-1 or VCAM-1 overnight at 4°C. Activated CD4^+^ T cells were serum starved for 3 hours in DMEM lacking all supplements, washed and resuspended in L15 media supplemented with 2 mg/mL D-glucose, and incubated 10 min at 37°C. 8.5 × 10^6^ T cells were then allowed to interact with surfaces coated with ICAM-1 or VCAM-1 for 20 mins at 37°C. Cell stimulated on surface-bound ICAM-1 or VCAM-1 were lysed by aspirating the media and adding 500 μL 1x ice cold lysis buffer (final composition: 1% Triton X-100, 150mM NaCl, 50mM Tris pH 7.5, 10mM MgCl_2_, 5mM NaF, 1mM sodium orthovanadate, and Roche EDTA-free protease inhibitor cocktail). Unstimulated control cells were lysed in solution by adding an equal volume of 2x cold lysis buffer for a final volume of 500 μL. Lysates were incubated on ice with periodic vortexing for 15 min, followed by centrifugation for 10 min at 16,000g at 4°C. A 50 μL aliquot of each whole cell lysate was retained, mixed with 4x sample buffer containing DTT (50mM final), and heated to 95°C for 10 min prior to separation by SDS-PAGE. The remainder of each lysate was used for immunoprecipitation.

For immunoprecipitation, 60 μL of Protein A agarose bead slurry (Repligen) were washed and resuspended in lysis buffer. Beads were then pre-bound to anti-phosphotyrosine (PY-20, 2.5 μg per condition) overnight at 4°C, with rotation. Antibody-charged beads were then washed 3x times with lysis buffer, mixed with 450 μL cell lysates, and rotated at 4°C overnight. Beads were then washed 3x with lysis buffer, mixed with 30 μL of 2x sample buffer, boiled, and analyzed by SDS-PAGE.

### Western Blotting

Proteins were separated by SDS-PAGE using the Invitrogen Novex Mini-cell system with NuPAGE 4-12% BisTris gradient gels. Proteins were transferred to nitrocellulose membranes (0.45 μm, BioRad) and blocked using LI-COR blocking buffer mixed 1:1 with PBS for 1 hr at RT. Primary antibodies were mixed in TBS 0.1% tween-20 (TBST) with 2% BSA and incubated with membranes overnight at 4°C on a shaker. Membranes were then washed 3x for 10 min each in TBST, and incubated with fluorophore-conjugated secondary antibodies in TBST with 2% BSA for 1 hr at RT. Membranes were washed 3x for 10 min in TBST, and imaged using a LI-COR Odyssey imaging system. Quantification of bands was done using ImageStudio software (LI-COR), and all measurements were within the linear range.

### Statistical analysis

Statistics were calculated and graphs were prepared using GraphPad Prism 8. When only two groups were being compared, a t-test was used. When more than two groups were compared, a one-way ANOVA was performed using multiple comparisons with a Tukey correction. *p<0.05; **p<0.01; ***p<0.001

## Acknowledgments

We thank the CHOP Flow Cytometry Core for access to flow cytometers. We also thank members of the Burkhardt and Hammer labs for discussions and critical reading of the manuscript.

## REFERENCES

1. Nourshargh S, Alon R. 2014. Leukocyte migration into inflamed tissues. Immunity 41:694–707.

2. Denucci CC, Mitchell JS, Shimizu Y. 2009. Integrin function in T-cell homing to lymphoid and nonlymphoid sites: getting there and staying there. Crit Rev Immunol 29:87–109.

3. Ley K, Laudanna C, Cybulsky MI, Nourshargh S. 2007. Getting to the site of inflammation: the leukocyte adhesion cascade updated. Nat Rev Immunol 7:678–689.

4. Gorina R, Lyck R, Vestweber D, Engelhardt B. 2014. beta2 integrin-mediated crawling on endothelial ICAM-1 and ICAM-2 is a prerequisite for transcellular neutrophil diapedesis across the inflamed blood-brain barrier. J Immunol 192:324–337.

5. Phillipson M, Heit B, Colarusso P, Liu L, Ballantyne CM, Kubes P. 2006. Intraluminal crawling of neutrophils to emigration sites: a molecularly distinct process from adhesion in the recruitment cascade. J Exp Med 203:2569–2575.

6. Schenkel AR, Mamdouh Z, Muller WA. 2004. Locomotion of monocytes on endothelium is a critical step during extravasation. Nat Immunol 5:393–400.

7. Shaw SK, Bamba PS, Perkins BN, Luscinskas FW. 2001. Real-time imaging of vascular endothelial-cadherin during leukocyte transmigration across endothelium. J Immunol 167:2323–2330.

8. Sumagin R, Sarelius IH. 2010. Intercellular adhesion molecule-1 enrichment near tricellular endothelial junctions is preferentially associated with leukocyte transmigration and signals for reorganization of these junctions to accommodate leukocyte passage. J Immunol 184:5242–5252.

9. Phillipson M, Heit B, Parsons SA, Petri B, Mullaly SC, Colarusso P, Gower RM, Neely G, Simon SI, Kubes P. 2009. Vav1 is essential for mechanotactic crawling and migration of neutrophils out of the inflamed microvasculature. J Immunol 182:6870–6878.

10. Bartholomaus I, Kawakami N, Odoardi F, Schlager C, Miljkovic D, Ellwart JW, Klinkert WE, Flugel-Koch C, Issekutz TB, Wekerle H, Flugel A. 2009. Effector T cell interactions with meningeal vascular structures in nascent autoimmune CNS lesions. Nature 462:94–98.

11. Steiner O, Coisne C, Cecchelli R, Boscacci R, Deutsch U, Engelhardt B, Lyck R. 2010. Differential roles for endothelial ICAM-1, ICAM-2, and VCAM-1 in shear-resistant T cell arrest, polarization, and directed crawling on blood-brain barrier endothelium. J Immunol 185:4846–4855.

12. Anderson NR, Buffone A, Jr., Hammer DA. 2019. T lymphocytes migrate upstream after completing the leukocyte adhesion cascade. Cell Adh Migr 13:163–168.

13. Valignat MP, Theodoly O, Gucciardi A, Hogg N, Lellouch AC. 2013. T lymphocytes orient against the direction of fluid flow during LFA-1-mediated migration. Biophys J 104:322–331.

14. Valignat MP, Negre P, Cadra S, Lellouch AC, Gallet F, Henon S, Theodoly O. 2014. Lymphocytes can self-steer passively with wind vane uropods. Nat Commun 5:5213.

15. Hornung A, Sbarrato T, Garcia-Seyda N, Aoun L, Luo X, Biarnes-Pelicot M, Theodoly O, Valignat MP. 2020. A Bistable Mechanism Mediated by Integrins Controls Mechanotaxis of Leukocytes. Biophys J 118:565–577.

16. Dominguez GA, Anderson NR, Hammer DA. 2015. The direction of migration of T-lymphocytes under flow depends upon which adhesion receptors are engaged. Integr Biol (Camb) 7:345–355.

17. Kim SHJ, Hammer DA. 2019. Integrin crosstalk allows CD4+ T lymphocytes to continue migrating in the upstream direction after flow. Integr Biol (Camb) 11:384–393.

18. Tedford K, Steiner M, Koshutin S, Richter K, Tech L, Eggers Y, Jansing I, Schilling K, Hauser AE, Korthals M, Fischer KD. 2017. The opposing forces of shear flow and sphingosine-1-phosphate control marginal zone B cell shuttling. Nat Commun 8:2261.

19. Buffone A, Jr., Anderson NR, Hammer DA. 2018. Migration against the direction of flow is LFA-1-dependent in human hematopoietic stem and progenitor cells. J Cell Sci 131.

20. Buffone A, Jr., Anderson NR, Hammer DA. 2019. Human Neutrophils Will Crawl Upstream on ICAM-1 If Mac-1 Is Blocked. Biophys J 117:1393–1404.

21. Kelleher D, Murphy A, Cullen D. 1990. Leukocyte function-associated antigen 1 (LFA-1) is a signaling molecule for cytoskeletal changes in a human T cell line. Eur J Immunol 20:2351–2354.

22. Porter JC, Bracke M, Smith A, Davies D, Hogg N. 2002. Signaling through integrin LFA-1 leads to filamentous actin polymerization and remodeling, resulting in enhanced T cell adhesion. J Immunol 168:6330–6335.

23. Smith A, Bracke M, Leitinger B, Porter JC, Hogg N. 2003. LFA-1-induced T cell migration on ICAM-1 involves regulation of MLCK-mediated attachment and ROCK-dependent detachment. J Cell Sci 116:3123–3133.

24. Roy NH, MacKay JL, Robertson TF, Hammer DA, Burkhardt JK. 2018. Crk adaptor proteins mediate actin-dependent T cell migration and mechanosensing induced by the integrin LFA-1. Sci Signal 11.

25. Sato T, Tachibana K, Nojima Y, D’Avirro N, Morimoto C. 1995. Role of the VLA-4 molecule in T cell costimulation. Identification of the tyrosine phosphorylation pattern induced by the ligation of VLA-4. J Immunol 155:2938–2947.

26. Hunter AJ, Shimizu Y. 1997. Alpha 4 beta 1 integrin-mediated tyrosine phosphorylation in human T cells: characterization of Crk- and Fyn-associated substrates (pp105, pp115, and human enhancer of filamentation-1) and integrin-dependent activation of p59fyn1. J Immunol 159:4806–4814.

27. Meng F, Lowell CA. 1998. A beta 1 integrin signaling pathway involving Src-family kinases, Cbl and PI-3 kinase is required for macrophage spreading and migration. EMBO J 17:4391–4403.

28. van Seventer GA, Mullen MM, van Seventer JM. 1998. Pyk2 is differentially regulated by beta1 integrin- and CD28-mediated co-stimulation in human CD4+ T lymphocytes. Eur J Immunol 28:3867–3877.

29. Tabassam FH, Umehara H, Huang JY, Gouda S, Kono T, Okazaki T, van Seventer JM, Domae N. 1999. Beta2-integrin, LFA-1, and TCR/CD3 synergistically induce tyrosine phosphorylation of focal adhesion kinase (pp125(FAK)) in PHA-activated T cells. Cell Immunol 193:179–184.

30. van Seventer GA, Salmen HJ, Law SF, O’Neill GM, Mullen MM, Franz AM, Kanner SB, Golemis EA, van Seventer JM. 2001. Focal adhesion kinase regulates beta1 integrin-dependent T cell migration through an HEF1 effector pathway. Eur J Immunol 31:1417–1427.

31. Cheung SM, Ostergaard HL. 2016. Pyk2 Controls Integrin-Dependent CTL Migration through Regulation of De-Adhesion. J Immunol 197:1945–1956.

32. Raab M, Lu Y, Kohler K, Smith X, Strebhardt K, Rudd CE. 2017. LFA-1 activates focal adhesion kinases FAK1/PYK2 to generate LAT-GRB2-SKAP1 complexes that terminate T-cell conjugate formation. Nat Commun 8:16001.

33. Huang B, Johansen KH, Schwartzberg PL. 2019. Efficient CRISPR/Cas9-Mediated Mutagenesis in Primary Murine T Lymphocytes. Curr Protoc Immunol 124:e62.

34. Schaff UY, Yamayoshi I, Tse T, Griffin D, Kibathi L, Simon SI. 2008. Calcium flux in neutrophils synchronizes beta2 integrin adhesive and signaling events that guide inflammatory recruitment. Ann Biomed Eng 36:632–646.

35. Dixit N, Yamayoshi I, Nazarian A, Simon SI. 2011. Migrational guidance of neutrophils is mechanotransduced via high-affinity LFA-1 and calcium flux. J Immunol 187:472–481.

36. Dixit N, Kim MH, Rossaint J, Yamayoshi I, Zarbock A, Simon SI. 2012. Leukocyte function antigen-1, kindlin-3, and calcium flux orchestrate neutrophil recruitment during inflammation. J Immunol 189:5954–5964.

37. Abram CL, Lowell CA. 2009. The ins and outs of leukocyte integrin signaling. Annu Rev Immunol 27:339–362.

38. Schmidt MHH, Dikic I. 2005. The Cbl interactome and its functions. Nat Rev Mol Cell Biol 6:907–918.

39. Murphy MA, Schnall RG, Venter DJ, Barnett L, Bertoncello I, Thien CB, Langdon WY, Bowtell DD. 1998. Tissue hyperplasia and enhanced T-cell signalling via ZAP-70 in c-Cbl-deficient mice. Mol Cell Biol 18:4872–4882.

40. Naramura M, Kole HK, Hu RJ, Gu H. 1998. Altered thymic positive selection and intracellular signals in Cbl-deficient mice. Proc Natl Acad Sci U S A 95:15547–15552.

41. Thien CB, Bowtell DD, Langdon WY. 1999. Perturbed regulation of ZAP-70 and sustained tyrosine phosphorylation of LAT and SLP-76 in c-Cbl-deficient thymocytes. J Immunol 162:7133–7139.

42. Rao N, Lupher ML, Jr., Ota S, Reedquist KA, Druker BJ, Band H. 2000. The linker phosphorylation site Tyr292 mediates the negative regulatory effect of Cbl on ZAP-70 in T cells. J Immunol 164:4616–4626.

43. Thien CB, Blystad FD, Zhan Y, Lew AM, Voigt V, Andoniou CE, Langdon WY. 2005. Loss of c-Cbl RING finger function results in high-intensity TCR signaling and thymic deletion. EMBO J 24:3807–3819.

44. Balagopalan L, Barr VA, Sommers CL, Barda-Saad M, Goyal A, Isakowitz MS, Samelson LE. 2007. c-Cbl-mediated regulation of LAT-nucleated signaling complexes. Mol Cell Biol 27:8622–8636.

45. Riedl J, Flynn KC, Raducanu A, Gartner F, Beck G, Bosl M, Bradke F, Massberg S, Aszodi A, Sixt M, Wedlich-Soldner R. 2010. Lifeact mice for studying F-actin dynamics. Nat Methods 7:168–169.

46. Park TJ, Curran T. 2008. Crk and Crk-like play essential overlapping roles downstream of disabled-1 in the Reelin pathway. J Neurosci 28:13551–13562.

47. Huang Y, Clarke F, Karimi M, Roy NH, Williamson EK, Okumura M, Mochizuki K, Chen EJ, Park TJ, Debes GF, Zhang Y, Curran T, Kambayashi T, Burkhardt JK. 2015. CRK proteins selectively regulate T cell migration into inflamed tissues. J Clin Invest 125:1019–1032.

